# Activation of the urotensin-II receptor by anti-COVID-19 drug remdesivir induces cardiomyocyte dysfunction

**DOI:** 10.1101/2022.08.08.503256

**Authors:** Akiko Ogawa, Seiya Ohira, Tatsuya Ikuta, Yuri Kato, Shota Yanagida, Yukina Ishii, Yasunari Kanda, Motohiro Nishida, Asuka Inoue, Fan-Yan Wei

**Affiliations:** Department of Modomics Biology and Medicine, Institute of Development, Aging and Cancer (IDAC), Tohoku University; Sendai, Miyagi, 980-8575, Japan.; Laboratory of Molecular & Cellular Biochemistry, Graduate School of Pharmaceutical Sciences, Tohoku University, 6-3, Aoba, Aramaki, Aoba-ku, Sendai, Miyagi 980-8578, Japan.; Department of Physiology, Graduate School of Pharmaceutical Sciences, Kyushu University, Fukuoka, Japan.; Division of Pharmacology, National Institute of Health Sciences, Kanagawa 210-9501, Japan.; Division of Pharmaceutical Sciences, Graduate School of Medicine, Dentistry and Pharmaceutical Sciences, Okayama University, Okayama 700-8530, Japan.; National Institute for Physiological Sciences and Exploratory Research Center on Life and Living Systems, National Institutes of Natural Sciences, Okazaki, Japan.

## Abstract

Remdesivir is an antiviral drug used for COVID-19 treatment worldwide. Cardiovascular (CV) side effects have been associated with remdesivir; however, the underlying molecular mechanism remains unknown. Here, we performed a large-scale G-protein-coupled receptor (GPCR) screening in combination with structural modeling and found that remdesivir is a selective agonist for urotensin-II receptor (UTS2R). Functionally, remdesivir treatment induced prolonged field potential in human induced pluripotent stem cell (iPS)-derived cardiomyocytes and reduced contractility in neonatal rat cardiomyocytes, both of which mirror the clinical pathology. Importantly, remdesivir-mediated cardiac malfunctions were effectively attenuated by antagonizing UTS2R signaling. Finally, we characterized the effect of 110 single-nucleotide variants (SNVs) in *UTS2R* gene reported in genome database and found four missense variants that show gain-of-function effects in the receptor sensitivity to remdesivir. Collectively, our study illuminates a previously unknown mechanism underlying remdesivir-related CV events and that genetic variations of *UTS2R* gene can be a potential risk factor for CV events during remdesivir treatment, which collectively paves the way for a therapeutic opportunity to prevent such events in the future.

**One Sentence Summary:** Remdesivir‘s activity as a selective agonist of urotensin-II receptor underlies its known cardiotoxicity in anti-viral therapy.

## INTRODUCTION

Nucleoside analogs have a long history in the field of drug design for antiviral treatment because nucleosides are used as building blocks for both DNA and RNA synthesis during viral replication (*1*). The primary mechanism of nucleoside analogs’ antiviral activities is attributed to inhibition of the viral RNA-dependent RNA polymerase (RdRp) (*2*). In response to the global pandemic of COVID-19, several nucleoside analogues, such as remdesivir, molnupiravir, and favipiravir, have been developed to treat the disease.

Remdesivir (GS-5734; Veklury) is a modified adenosine analogue which contains a McGuigan prodrug moiety including phenol and L-alanine ethylbutyl ester which increases its lipophilicity and cell permeability (*3*). Following intravenous administration, remdesivir is rapidly converted into the mono-nucleoside form (GS-441524) and is intracellularly metabolized by multiple host enzymes to its pharmacologically active triphosphate form, which, in turn, acts as a potent and selective inhibitor of the RNA-dependent RNA polymerase of multiple viruses (*4*) (*5*). Remdesivir was initially used for the treatment of Ebola virus (*6*) and has been approved for the treatment of Coronavirus Disease 2019 (COVID-19) amid the global pandemic. Remdesivir shortened the time to recovery in adults who were hospitalized with COVID-19 and had evidence of lower respiratory tract infection (*7*). More recent data demonstrated a significant reduction in hospitalizations with a 3-day course of intravenous remdesivir (*8*). Although remdesivir is generally well tolerated by most individuals, common adverse events for remdesivir have been reported, including rash, headache, nausea, diarrhea, and elevated transaminases (*7*). Current guidelines for remdesivir suggest a careful monitoring of liver function during treatment and recommend against its use in patients with renal dysfunction (*9*). Additionally, cardiovascular events, including hypotension, bradycardia, QT prolongation and T-wave abnormality have been reported (*10–13*). When administrated intravenously, remdesivir shows broad tissue distribution, including to the heart (*14*), but the precise molecular mechanism underlying the cardiovascular side-effects of remdesivir remains unclear.

Molnupiravir (EIDD-2801/MK-4482; Lagevrio) has been authorized for emergency use by the FDA under an Emergency Use Authorization (EUA) for the treatment of mild-to-moderate COVID-19 in adults who are at high-risk for progression to severe COVID-19. Molnupiravir is an oral cytosine analogue which contains an isopropylester prodrug of the β-D-N^4^-hydroxycytidine (NHC). The active form of NHC is a substrate of RdRp and impairs the fidelity of SARS-CoV-2 replication, provoking “error catastrophe” (*15*). A clinical trial in nonhospitalized adults showed that early treatment with molnupiravir effectively reduced the risk of hospitalization or death in at-risk, unvaccinated adults with COVID-19 (*16*). In addition to molnupiravir, favipiravir (T-705; Avigan), which is an anti-influenza drug, has been under clinical trial for the treatment of COVID-19. Favipiravir s a nucleobase analogue derived from pyrazine carboxamide (6-fluoro-3-hydroxy- 2-pyrazinecarboxamide) (*17*). The suggested modes of action of favipiravir comprises a mix of both chain termination and mutator events (*18*). Several adverse events have been reported during use of molnupiravir and favipiravir, including diarrhea, dizziness, and nausea for molnupiravir (*19*), and hyperuricemia and increased alanine aminotransferase for favipiravir (*20*). Importantly, unlike remdesivir, cardiovascular side effects have not been reported with the use of molnupiravir or favipiravir.

In addition to their function as building blocks for DNA/RNA synthesis, nucleotides/nucleosides can act as endogenous ligands for G-protein-coupled receptors (GPCR) and induce diverse pathophysiological responses (*21–25*). Given the presence of nucleoside- mimicking structures in remdesivir, molnupiravir, and favipiravir, we hypothesized that these drugs may cause side effects by directly activating GPCR. Here, we report that remdesivir, but not molnupiravir and favipiravir, is a selective ligand for the urotensin-II receptor (UTS2R) and causes cardiac dysfunction.

## RESULTS

### Remdesivir is a selective agonist of the urotensin-II receptor (UTS2R)

We screened anti COVID-19 drugs including remdesivir, molnupiravir, and favipiravir, against 348 GPCRs using a transforming growth factor-α (TGF-α) shedding assay (*26*). We used chimeric Gα subunit proteins for the initial screening to efficiently detect the receptor activation regardless of the type of Gα subunit involved (*26*). Among the three drugs, we discovered that remdesivir is a selective activator for the urotensin-II receptor (UTS2R) (Fig. 1A, Data set S1). As remdesivir potently induced UTS2R response without any chimeric Gα protein, the subsequent UTS2R analysis were performed without the exogenous addition of the chimeric Gα protein. A concentration-response analysis revealed that the half-maximal effective concentration (pEC_50_) of remdesivir was 4.89 ± 0.03 (EC_50_ = 13 μM, Fig. 1B left). It should be noted that the potency of remdesivir toward UTS2R is lower than that of endogenous peptidic ligand urotensin-II (UT2, pEC_50_ = 10.72 ± 0.04; EC_50_ = 21 fM, Fig. 1B right). Nevertheless, remdesivir administration at a clinical dosage is considered to be within the working range of agonistic effects because the maximum plasma concentration of remdesivir reaches to 9.03 μM after intravenous injection in healthy adults (*27*). Interestingly, unlike UT2, remdesivir failed to induce a β-arrestin recruitment response and thus behaved as a G-protein-biased ligand (Fig. 1C, S1A).

**Fig. 1.**
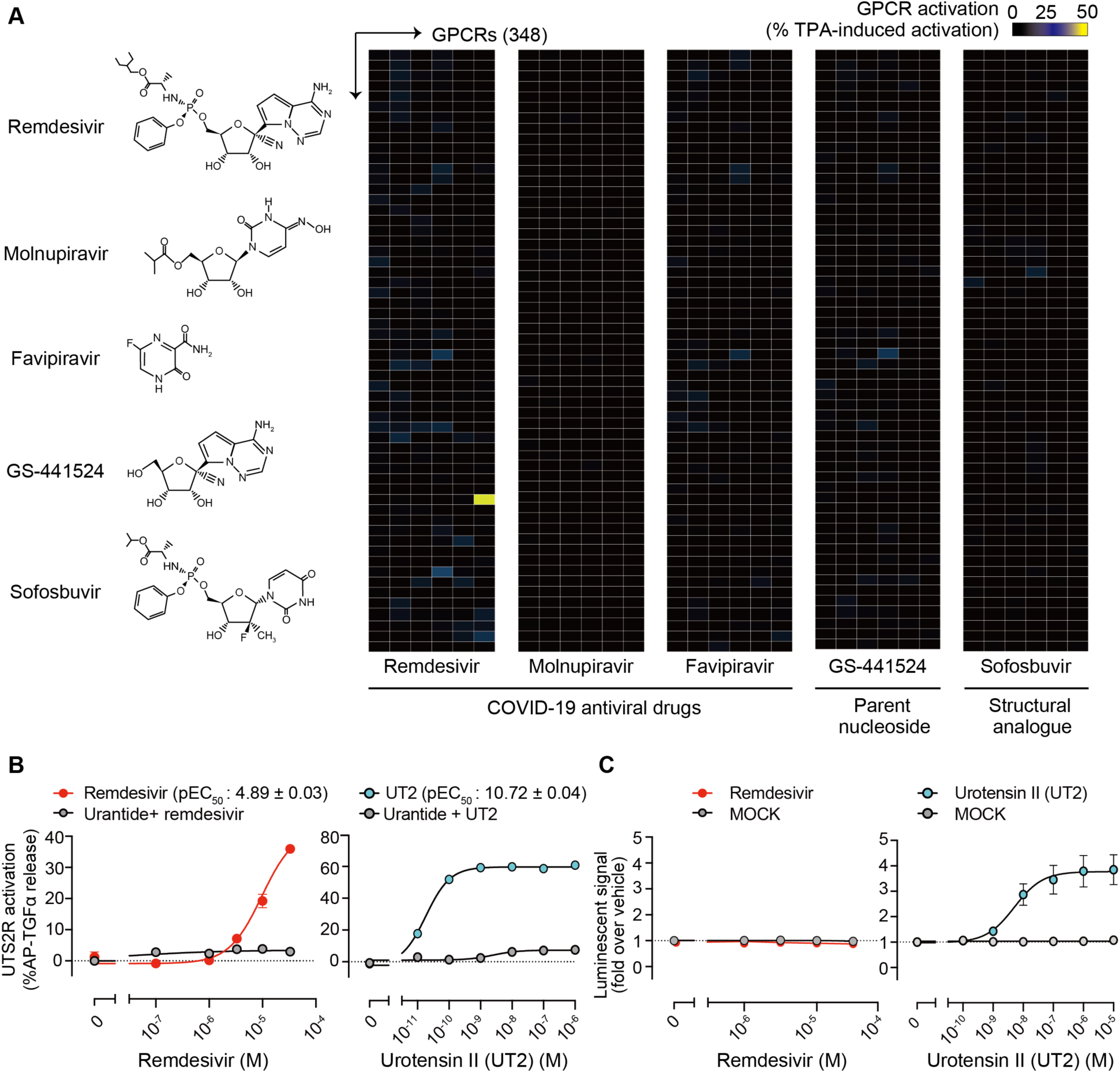
Identification of remdesivir as a selective agonist of urotensin-II receptor (UTS2R). **(A)** Left, Chemical structures of tested compounds. Right, Heatmap showing the relative levels of GPCR activation as measured by the TGFα-shedding assay. Color scale represents the % GPCR activation compared to TPA (12-O-tetradecanoylphorbol 13-acetate)- mediated receptor activation, which induces the maximum TGFα-shedding response independently of GPCR. Yellow cell represents activation of UTS2R by remdesivir. The tested compounds’ concentrations were 10 μM for remdesivir, molnupiravir, GS-441524, and sofosbuvir, and 100 μM for favipiravir. **(B)** TGFα-shedding response curves for UTS2R by remdesivir (left) and urotensin-II (UT2, right) in the presence or absence of urantide, a UTS2R antagonist. The half-maximal effective concentration (pEC_50_) of remdesivir was 4.89 ± 0.03 (EC_50_ = 13 μM, E_max_ = 47 ± 1.4 %AP-TGFα release). The half- maximal effective concentration (pEC_50_) of UT2 was 10.72 ± 0.04 (EC_50_ = 21 fM, E_max_ = 59 ± 0.70 %AP-TGFα release). **(C)** Remdesivir (left)- or urotensin-II (UT2, right)- mediated β-arrestin 1 recruitment assay for UTS2R. Data are shown as means ± SEM (n≥3).

We then investigated whether the metabolites of remdesivir activate UTS2R. GS-441524 and GS-704277, the major and minor metabolites of remdesivir (*6, 27*), respectively, showed no effect on UTS2R activation (Figs. 1A, S1B). To investigate whether the McGuigan prodrug moiety is responsible for remdesivir’s UTS2R activation, we tested sofosbuvir (GS-7977), an FDA-approved McGuigan-class prodrug (*3*), against UTS2R. However, sofosbuvir did not activate UTS2R (Figs. 1A, S1B). These results suggest that both the McGuigan prodrug moiety and nucleoside base of remdesivir are required to activate the receptor.

### Molecular basis of UTS2R activation by remdesivir

UTS2R belongs to the class A GPCR family (*28*) and consists of the canonical 7 transmembrane helices (TM), the amphipathic helix 8 at the C-terminus (H8), two antiparallel β- strands in the extracellular loop 2 (ECL2), a relatively short N-terminal domain with two N-glycosylation sites, and a palmitoylation anchor at the C-terminal tail (Fig. 2A upper). Meanwhile, remdesivir is an analogue of adenosine and a phosphoramidate prodrug of the McGuigan class (phenol and L-alanine ethylbutyl ester), which masks the anionic phosphate moiety on remdesivir, thus improving drug delivery. To further elucidate the molecular basis for ligand-receptor binding, we performed *in silico* structural docking of UTS2R in the presence of remdesivir (Fig. 2A lower). Our analysis showed multiple amino acid residues in the orthosteric pocket that potentially stabilize binding to remdesivir. First, the cyano group of the nucleosugar of remdesivir forms a hydrogen bond to the UTS2R residue T304^7.42x41^ (superscripts denote the generic GPCR numbering system (*29*)). Second, the phenyl group of the McGuigan prodrug moiety of remdesivir is capped with N297^7.35x34^. Finally, the amino group of the nucleobase of remdesivir forms a hydrogen bond with M134^3.36^ at the bottom of the pocket (NH···S hydrogen bond (*30*)).

**Fig. 2.**
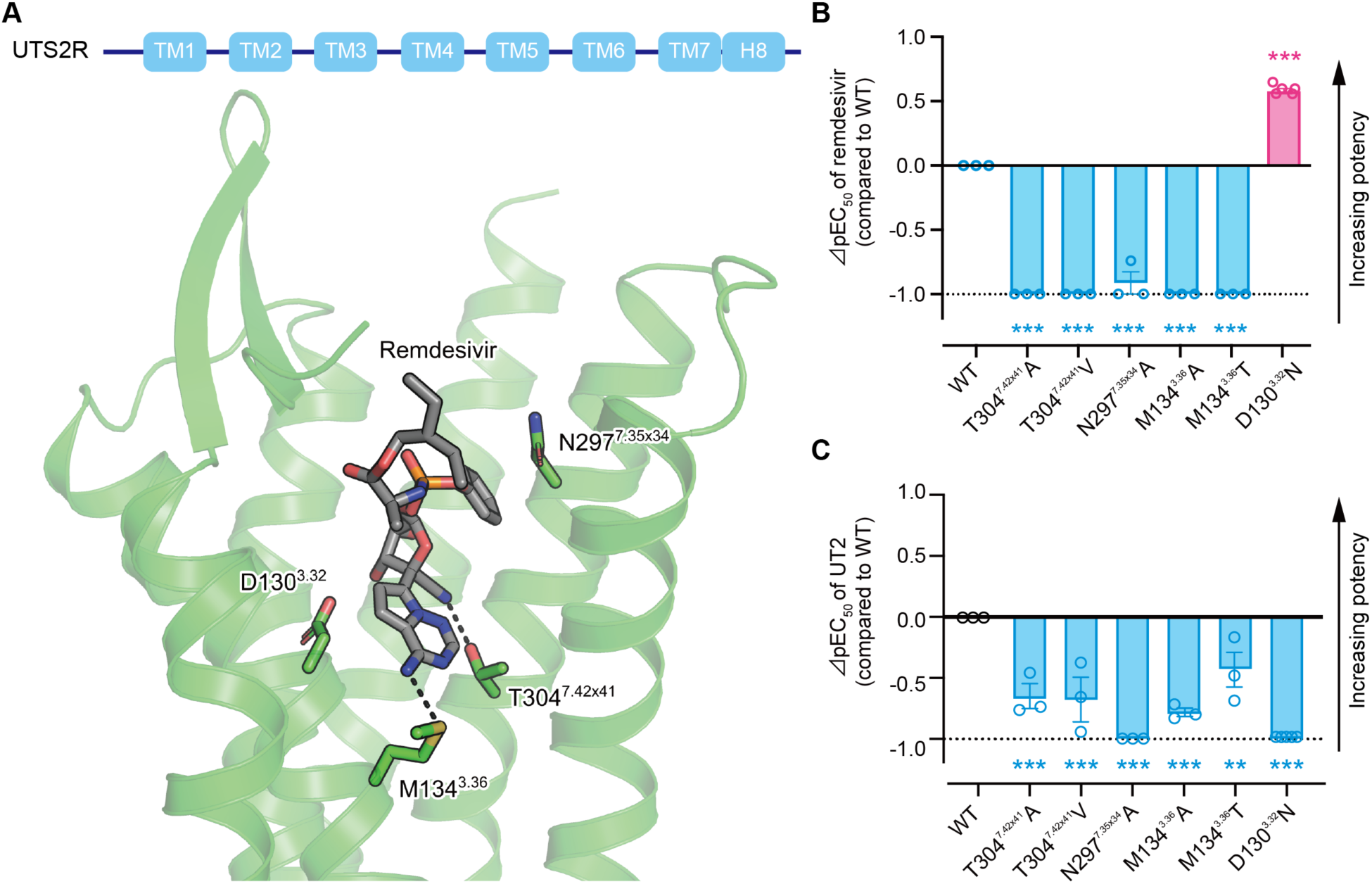
Docking simulation of UTS2R with remdesivir. **(A)** Upper panel, Schematic of UTS2R; Lower panel, Docking model of UTS2R with remdesivir. AlphaFold structure of human UTS2R is shown as green ribbons, omitting TM5 for clarity. Remdesivir and selected side chains of the receptor are shown as sticks and colored gray and green, respectively. Black dashed lines indicate hydrogen bonds. **(B, C)** Effects of the indicated mutations on remdesivir (B)- or UT2 (C)- mediated receptor activation. The Y-axis (⊿pEC_50_ value) represents the relative activation potency of each mutant receptor compared to WT receptor. ⊿pEC_50_ value = pEC_50_ mutant - pEC_50_ WT. The ⊿pEC_50_ cutoff value was set to -1, as indicated by dashed lines. EC_50_ values were determined by the TGFα-shedding assay. ** p < 0.01, *** p < 0.001 vs WT by one-way ANOVA followed by Dunnett’s multiple comparison test. All data are shown as means ± SEM (n ≥3).

To validate the docking model, we mutated these UTS2R residues (T304^7.42x41^, N297^7.35x34^, and M134^3.36^) to other amino acids and tested whether these mutations affect remdesivir-mediated UTS2R activation (Figs. 2B,C Fig. S2A, B). In line with the *in silico* simulation, mutation of the three residues almost completely abolished the activation potency of remdesivir to UTS2R, indicating that remdesivir needs to contact to UTS2R at multiple residues to achieve stable binding. Both nucleoside and McGuigan moiety are essential for receptor binding, with N297^7.35x34^ interacting with the McGuigan moiety and M134^3.36^ and T304^7.42x41^ interacting with the nucleoside moiety. This structural observation explains why mutating a single amino acid strikingly reduces the remdesivir’s activity towards UTS2R. This explanation is further supported by the result that UTS2R does not respond to any remdesivir metabolites, from which the McGuigan moiety is metabolically removed. In contrast, these mutations only partially reduced the UT2-mediated UTS2R activation. Meanwhile, contrary to these three residues, we found that D130^3.32^ has an inhibitory effect on remdesivir-UTS2R binding. The D130^3.32^N mutation abolished UT2-mediated UTS2R activation but significantly upregulated remdesivir-mediated receptor activation (Figs. 2B, C Fig. S2A, B). According to the docking model, the D130^3.32^ residue is localized near the nucleobase moiety of remdesivir (Fig. 2A). We speculate that the negative charge of D130^3.32^ residue may induce an electron repulsion effect (*31*), leading to destabilization of the interaction between D130^3.32^ residue and the nucleobase. These results demonstrate that remdesivir-mediated UTS2R activation is mediated by specific binding between UTS2R and its McGuigan prodrug moiety and nucleoside base, which differs from that of UTS2R and the endogenous ligand.

### Remdesivir-mediated UTS2R activation underlies drug-mediated cardiotoxicity

To determine whether remdesivir-mediated UTS2R activation induces intracellular signaling transduction, we stimulated UTS2R-expressing HEK293 cells with remdesivir and examined the phosphorylation status of extracellular signal-regulated kinase (ERK)1/2. Application of remdesivir for up to 72 h evoked long-lasting and dose-dependent phosphorylation of ERK1/2 (Figs. 3A, S3A). Importantly, remdesivir-mediated ERK phosphorylation was abolished by the UTS2R antagonist (Figs. 3A, S3A). The response of ERK1/2 induced by remdesivir was similar to that induced by UT2 (Fig. S3B).

**Fig. 3.**
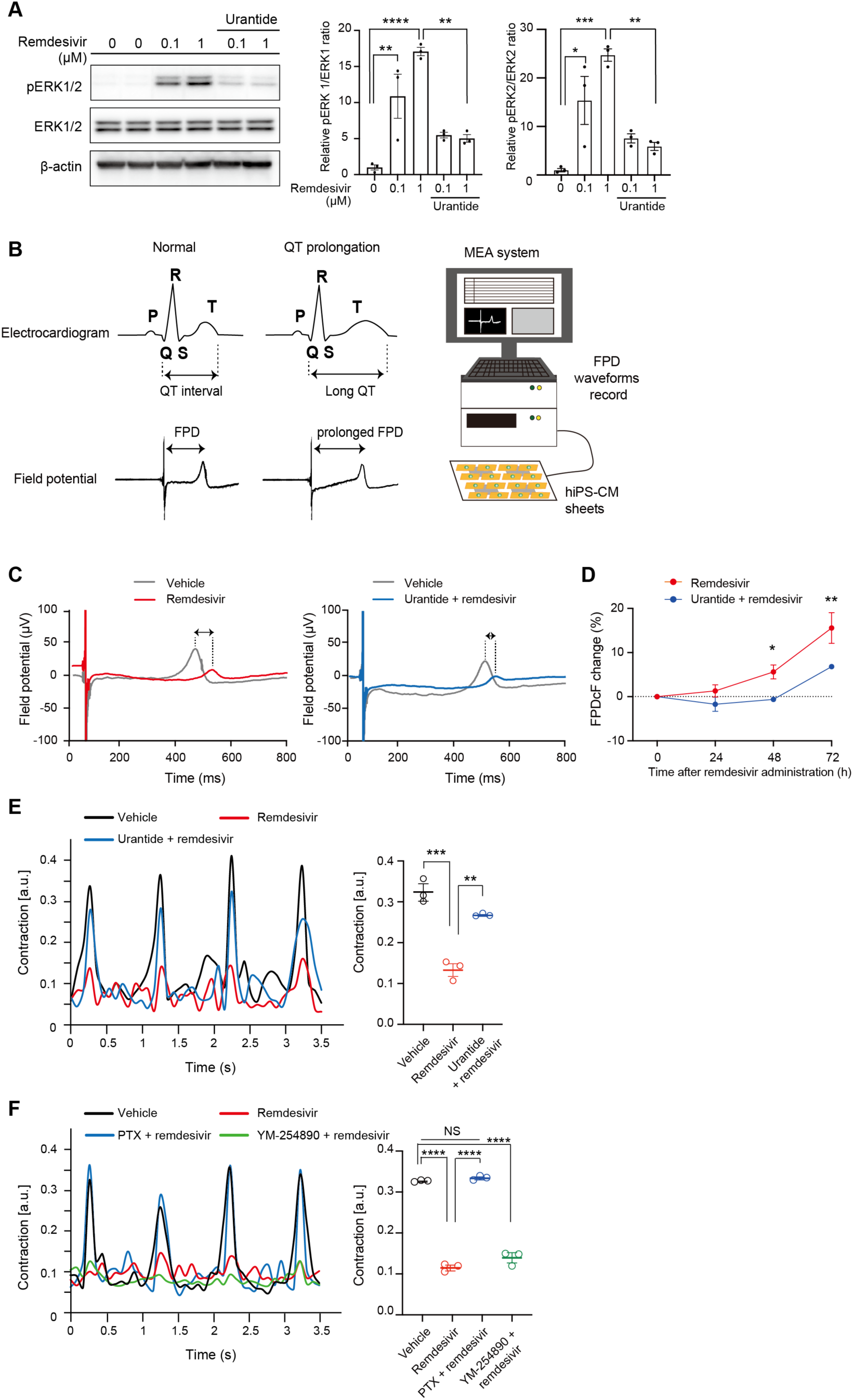
Cardiotoxic effects of remdesivir-mediated UTS2R activation. **(A)** Serum-starved HEK293 cells overexpressing UTS2R were stimulated with the indicated concentrations of remdesivir for 5 min with or without urantide, a UTS2R antagonist, and the lysates subjected to western blotting analysis. ERK1 and ERK2 activation ratio (pERK1/ERK1 and pERK2/ERK2) were calculated with data normalized to the vehicle. * p <0.05, ** p <0.01, *** p <0.001, **** p <0.0001 by Tukey’s multiple comparisons test. **(B)** Left, Temporal correlation between the field potential duration and the QT interval on the surface ECG; Right, Schematic of multielectrode array (MEA) platform. **(C)** Representative field potential waveform in human induced pluripotent stem cell-derived cardiomyocytes (hiPSC-CMs) treated with 1 µM remdesivir in the presence or absence of urantide for 72 h. **(D)** Effect of remdesivir and urantide on field potential prolongation in hiPSC-CMs. * p <0.05, ** p <0.01 by two-way ANOVA followed by Šídák’s multiple comparisons test. **(E-F)** Left, Representative waveform of contractility under pacing in neonatal rat cardiomyocytes (NRCMs) by 1 µM remdesivir with or without (E) urantide or (F) pertussis toxin (PTX), a Gα_i/o_ inhibitor, and YM-254890, a Gα_q/11_ inhibitor, at 48 h. Right, Effect of remdesivir with (E) urantide or (F) PTX and YM-254890 on contractility under pacing in NRCMs. ** p <0.01, *** p <0.001, **** p <0.0001 by Tukey’s multiple comparisons test. All data in Fig. 3 are represented as means ± SEM (n ≥3).

UT2 and its receptor UTS2R are widely expressed in tissues, with relatively high expression in cardiovascular systems (*32*) (Fig. S3C, D). Prompted by the cardiotoxic potential of remdesivir (*10–13*), we assessed the impact of remdesivir on cardiomyocyte functions. We examined the effect of remdesivir on the field potential (FP) using human-induced pluripotent stem cell-derived cardiomyocytes (hiPSC-CMs) (*33*), in which the expression level of *UTS2R* is comparable to that of the human heart (Fig. S3E). The field potential duration (FPD) correlates closely with the QT interval on an electrocardiogram (ECG) (*34*) (Fig. 3B left). Notably, the prolongation of the QT interval has been linked with the occurrence of severe and potentially fatal arrhythmias, and QT prolongation is the leading cause of drug-induced cardiovascular toxicity (*35*). For the assessment of FPD, the use of a multielectrode array (MEA) platform is well-accepted for its ability to monitor electrophysiology of cardiomyocytes at the cell population level (Fig. 3B right) (*34*). An MEA analysis revealed that hiPSC-CMs treated with remdesivir showed a prolonged FPD by 1.32 ± 1.38% at 24 h, 5.60 ± 1.60% at 48 h, and 15.57 ± 3.49% at 72 h, respectively. Notably, the prolongation was significantly suppressed by UTS2R antagonist (Fig. 3C, D). Indeed, application of UTS2R antagonist with remdesivir showed almost no FPD delay at 24 h (-1.71 ± 1.60 %) and 48 h (-0.61 ± 0.11 %), while the reversal effect was still significant but became partial at 72 h (Fig. 3D). These results elucidate a previously unknown mechanism of the reported proarrhythmic risks of remdesivir (*11–13*), which is at least partially dependent on UTS2R. Next, we assessed the effects of remdesivir on cardiac contractility. To this end, we used neonatal rat cardiomyocytes (NRCMs) and evaluated the contraction force. Under a constant pacing protocol, NRCMs with chronic remdesivir treatment showed reduced contractility, which was significantly attenuated by the UTS2R antagonist (Fig. 3E).

Heterotrimeric G proteins, including Gα_s_, Gα_i/o_, Gα_q/11_, and Gα_12/13_, are the downstream effectors of GPCRs. Among those, the Gα_i/o_ family has been implicated in the myocardial contractility and heart rate via the modulation of ion channels (*36*). Since UTS2R is coupled to Gα_i/o_ and Gα_q_ (*28*), we sought to determine which Gα protein is involved in the remdesivir-mediated decrease in myocardial contraction. Gα_i/o_ inhibitor pertussis toxin (PTX), but not Gα_q/11_ inhibitor YM-254890, completely blocked the effect of remdesivir and restored the peak contractions of NRCMs (Fig. 3F). A previous study demonstrated how activating Gα_i/o_ in NRCMs leads to liberation of Gβγ, which intracellularly activates PI3K, leading to the activation of AKT and ERK1/2 (*37*). In line with this observation, Gα_i/o_ inhibition reduced the remdesivir-induced phosphorylation of ERK1/2 and AKT (Fig. S3F). These results suggest that remdesivir reduces cardiomyocyte contractility through the Gα_i/o_-dependent PI3K/AKT/ERK signal transduction pathway. Collectively, our results indicate that remdesivir itself can function as an exogenous ligand of UTS2R. Moreover, we identified the proarrhythmic and negative inotropic potential of remdesivir, both of which are UTS2R dependent.

### Genetic effects of remdesivir-mediated UTS2R activation

To understand the impact of genetic variance on the susceptibility to remdesivir-UTS2R signaling in humans, we extract SNV information from a genome database that includes the 14KJPN Genome Reference Panel, which is a large-scale population-based genomic database constructed from the DNA sequence of 14,000 Japanese individuals (*38*), followed by functional assay on receptor activation. A total of 2178 variants are reported in the *UTS2R* locus, of which 139 variants are missense variants (Data set S2). A considerable number of missense SNVs listed in the 14KJPN are also reported in the gnomAD database (https://gnomad.broadinstitute.org), which contains SNVs from broader ethnicities.

Accordingly, we generated 110 missense mutants corresponding to the human SNVs in the *UTS2R* gene. We excluded 29 missense mutations in the 5’-region of *UTS2R* as the confidence score for the structural prediction of the N-terminus was relatively low. Among the 110 missense SNVs, 44 SNVs showed a decrease in the sensitivity to remdesivir compared to the WT receptor (⊿pEC_50_ <-0.3, which corresponds to over a twofold EC_50_ increase compared to WT receptor; Fig. 4A upper panel). Meanwhile, 47 SNVs displayed a decreased sensitivity to UT2 compared to WT receptor, of which 18 SNVs overlapped with those that showed a decreased sensitivity to remdesivir (Figs. 4A lower panel, S4A). Notably, we found four missense SNVs (G68^1.49^C, D130^3.32^G, V159^34.54^M, and A249^ICL3^G) that can increase the receptor sensitivity toward remdesivir compared to the WT receptor (⊿pEC_50_ >0.3, which corresponds to a <0.5-fold EC_50_ decrease compared to the WT receptor; Fig. 4A-C). Furthermore, among these four gain-of-function remdesivir-sensitive UTS2R SNVs, the G68^1.48^C and D130^3.32^G mutants conversely exhibited a decrease in the sensitivity toward UT2, while the V159^34.54^M and A249^ICL3^ mutants showed a moderate or insignificant increase in UT2 sensitivity (⊿pEC_50_ <0.3; Fig. 4C, S4A). Collectively, the results suggest that individuals with a G68^1.49^C, D130^3.32^G, V159^34.54^M, or A249^ICL3^G mutation in the *UTS2R* gene are sensitive to remdesivir, possibly making them more susceptible to UTS2R-mediated cardiotoxicity, although the allele frequencies of the gain-of-function variants are low (Fig. S4B).

**Fig. 4.**
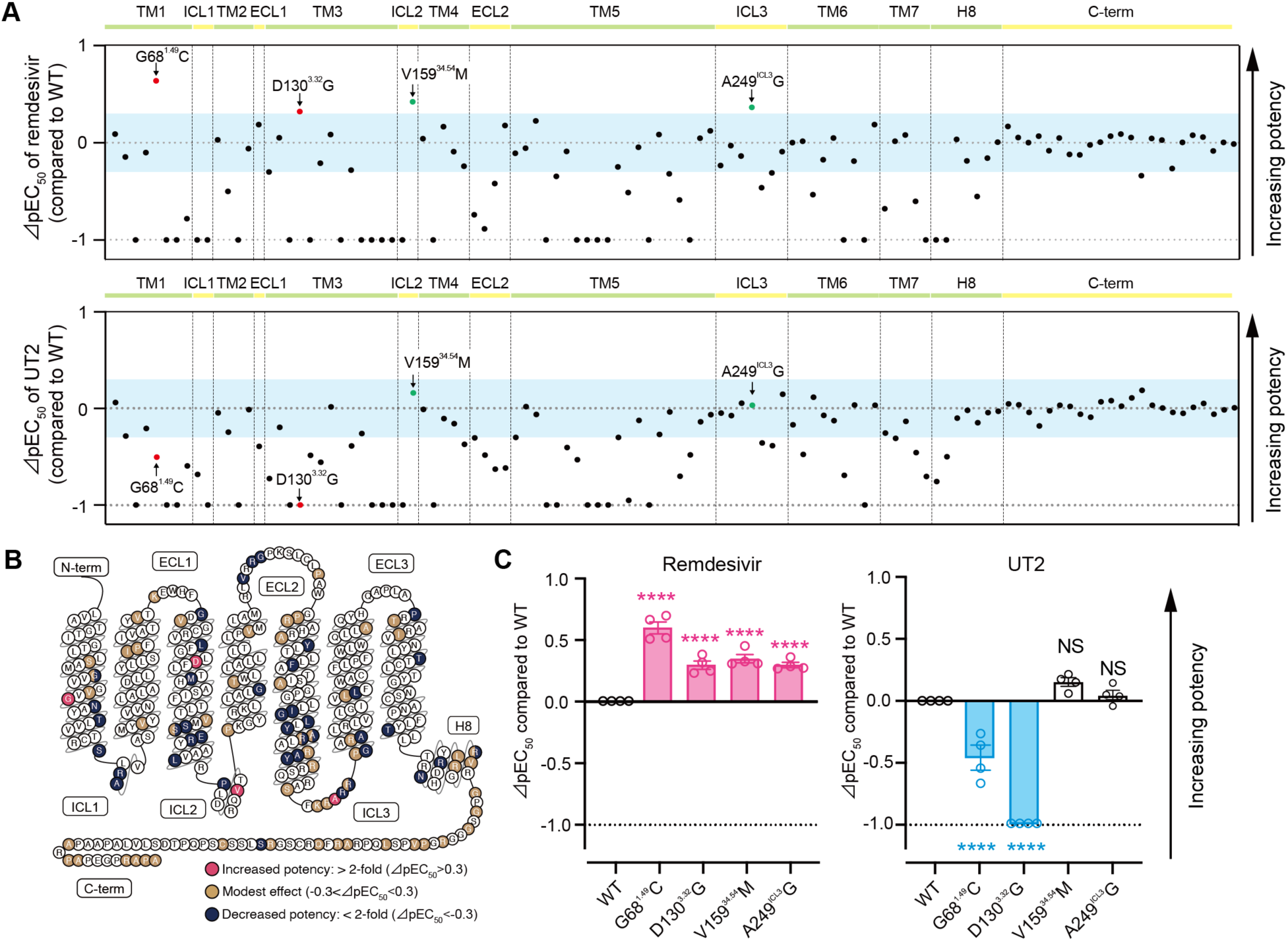
Genetic effects of remdesivir-mediated UTS2R activation. **(A)** Effects of 110 missense SNVs from the *UTS2R* gene on (upper panel) remdesivir-mediated and (lower panel) UT2- mediated receptor activation, measured by the TGFα-shedding assay. Y-axis (⊿pEC_50_ value) represents the relative activation potency of each mutant receptor compared to the WT receptor. The ⊿pEC_50_ cutoff value was set to -1, as indicated by dashed lines. Light blue bands represent the range of -0.3< ⊿pEC_50_ <0.3, which corresponds to the range of an 0.5-fold to 2-fold (less than 2-fold) change in the EC_50_ of the mutant receptor compared to the WT receptor. **(B)** Snake plot diagram of UTS2R showing locations of mutations and their effects on the potency of remdesivir (modified from www.gpcrdb.org). **(C)** Effects of the selected mutations on (left) remdesivir-mediated and (right) UT2-mediated receptor activation. EC50 values are determined by the TGFα-shedding assay. **** p <0.0001 vs. WT by one-way ANOVA followed by Dunnett’s multiple comparisons test. Data are shown as means ± SEM (n ≥3).

## DISCUSSION

Following the completion of the Adaptive COVID-19 Treatment Trial 1 (ACTT-1) (*7*), which demonstrated remdesivir’s superiority over placebo for improving time to recovery in hospitalized COVID-19 patients, remdesivir has been one of the most commonly prescribed medications for patients hospitalized for COVID-19 infection. Importantly, the ACTT-1 investigators reported that 0.2% of patients receiving remdesivir, but not patients receiving placebo, showed arrythmias (other than atrial fibrillation, supraventricular tachycardia, ventricular tachycardia, and ventricular fibrillation), although these cardiovascular effects was not considered as significant adverse effects. However, the percentage of patients experiencing arrythmias could be underestimated, since early clinical trials are insufficiently powered to detect uncommon adverse events (*39*). Indeed, a large retrospective pharmacovigilance cohort study, which used the medical records from more than 130 countries and 20 million patients, reported that cardiac arrest, bradycardia, and hypotension is associated with remdesivir use (*10*). The elevation of plasma remdesivir concentration is associated with an increase in field potential duration with decreased Na^+^ peak amplitudes and spontaneous beating rates, which might potentially induce prolonged QT interval and torsade de point (*40, 41*). Importantly, the cardiac adverse effects appear to be directedly caused by remdesivir, since they were reported to resolve within 24-48 hours of discontinuing remdesivir (*42, 43*). To date, the precise mechanism underlying the remdesivir’s cardiac side effects has remained unclear with no means of predicting the populations susceptible to its cardiac side effects and no specific treatment for the cardiotoxicity. Therefore, there is a pressing need to understand how remdesivir induces the cardiac dysfunction.

Using an unbiased and large-scale GPCR screening strategy, we identified that remdesivir, but not molnupiravir and favipiravir, selectively activates UTS2R. Interestingly, UTS2R is highly expressed in heart tissue, including cardiomyocytes. Activation of UTS2R by Urotensin-II (UT2), the endogenous ligand of UTS2R, has been implicated in cardiac dysfunction. For example, the plasma level of UT2 and expression level of UTS2R in cardiomyocytes are elevated in patients with end-stage congestive heart failure (*44*). In line with these previous studies, we found that activation of UTS2R by remdesivir at a concentration of 1 μM induced electrical abnormalities and contraction force impairment in cultured cardiomyocytes, both of which resemble the reported cardiac side effects in humans. Furthermore, these adverse effects were effectively blocked by antagonizing UTS2R or inhibiting its downstream signaling. Clinically, remdesivir is administrated intravenously at a dose of 200 mg once followed by 100 mg daily for a total of 5-10 days in adults and children ≥40 kg. The estimated peak plasma concentration of remdesivir is 9.03 μM in healthy adults and can be higher in patients with renal or hepatic impairment because of its renal and biliary excretion (*45*). Our results suggest that the clinical dosage of remdesivir is sufficient to activate UTS2R, and patients with renal or hepatic impairment may be at high risk for adverse events mediated through the remdesivir-UTS2R axis. Notably, our GPCR assay clearly showed that remdesivir has no effect on adenosine receptors, of which the activation can also impact cardiac functions. Thus, the activation of UTS2R is likely responsible for the cardiac side effects.

An important finding of this study is that single-nucleotide variants (SNVs) in the coding region of UTS2R have large impacts on its response to remdesivir. This result is in line with previous studies that show SNVs in GPCR genes are associated with the pathophysiology of various diseases, including CV diseases, and affect therapeutic outcomes (*46*). While 40% of SNVs in UTS2R (44/110) showed at least a twofold reduction in potency toward remdesivir compared to WT receptor, and 56% (62/110) showed no remarkable change (within the range of an 0.5-fold to 2-fold change), we identified four gain-of-function SNVs that showed more than two-fold increase in the response to remdesivir. Notably, D130^3.32^G was a variant at the same amino acid for which we identified a gain-of-function mutation (D130^3.32^N) using *in silico* modeling, indicating the importance of D130^3.32^ residue in remdesivir recognition.

Activation of GPCR often induces recruitment of β-arrestin to the receptor, followed by receptor internalization to the intracellular compartment. This internalization terminates GPCR activation and promotes secondary signaling pathways (*47*). Interestingly, unlike the endogenous ligand UT2, which effectively induces β-arrestin recruitment (Fig 1C) (*48*), remdesivir-mediated UTS2R activation did not induce β-arrestin recruitment. Thus, our results suggest that remdesivir is a G-protein-biased ligand (Fig. 1C, S1A). Such biased activation of UTS2R by remdesivir may have an important impact on cardiac function in a way that allows prolonged activation of UTS2R without β-arrestin-mediated shutdown and thereby an exaggeration of downstream signaling and enhanced cardiotoxic effects.

Upon ligand binding, GPCRs, including UTS2R, initiate conformational change that induces the activation of heterotrimeric G proteins and dissociation of Gα and Gβγ subunit complexes. Gα proteins include Gα_s_, Gα_i/o_, Gα_q/11_, and Gα_12/13_ proteins, which are responsible for downstream signaling transduction. Members of the Gα_i/o_ family are widely distributed, including in the cardiac system, where they are highly expressed and act to regulate myocardial contractility and heart rate via modulation of ion channels (*36*). For example, resting heart rate is controlled by cholinergic signals mediated through muscarinic M2 Gα_i_-coupled receptors. These effects occur by inhibition of adenylyl cyclase (AC) and by Gβγ-inhibition of a potassium channel in the sinoatrial node. The transduction mechanism of UTS2R is the coupling and activation of Gα_i/o_ as well as Gα_q_ (*28*), consistent with the UT2-UTS2R axis being implicated in CV regulation through complex signaling pathways, both physiologically and pathologically. Our results using NRCMs clearly showed that the remdesivir-mediated decrease in myocardial contraction is dependent on Gα_i/o_, but not Gα_q/11_ (Fig. 3F). Since NRCMs resemble the phenotype of atrial myocytes (*49*), the decrease in myocardial contractility under constant pacing by remdesivir indicates impairment of calcium handling (*50*), which is consistent with the slowing of the heart rate, a distinctive cardiovascular side effect of remdesivir (*10*).

Despite the discovery that remdesivir can activate UTS2R and cause cardiotoxicity, the lack of clinical evidence is a major limitation of this study. Yet, as the allele frequencies of the gain-of-function variants are low (Fig. S4B) and usage of remdesivir is expected to decline due to the recent prevalence of the Omicron COVID-19 variant with its milder symptoms, a large-scale clinical study on the association between remdesivir sensitivity and genomic variance is challenging to execute. Additionally, we have not determined the precise molecular mechanisms that induce the UTS2R-dependent proarrhythmic risk of remdesivir. Possible explanations are the impaired regulation of gene expression or trafficking of ERG potassium channels, which are essential for electrical activity in the heart (*51*). Alternatively, the chronic and cumulative cardiotoxic effects of remdesivir-UTS2R axis are consistent with a downstream impact on translation and transcription. In addition, a previous study suggested that remdesivir-related cardiotoxicity can be caused by mitochondrial dysfunction (*52*) since the active form of remdesivir shows an inhibitory effect toward mitochondrial RNA polymerase (mtRNAP) at a high dose (*53*). However, treatment with remdesivir at 10 μM, which is equivalent to the maximum plasma concentration following remdesivir administration in humans (*27*), did not affect the steady-state levels of mitochondrial respiratory complex proteins (Fig. S3G). Although we do not exclude the possibility that remdesivir might affect mitochondrial metabolism through UTS2R signal transduction, the current data suggests that the remdesivir-UTS2R axis is the major off-target of remdesivir.

In conclusion, to our knowledge, this is the first report showing that remdesivir is a selective agonist of UTS2R and that remdesivir-mediated UTS2R activation underlies drug-mediated cardiotoxicity. Furthermore, we discovered that specific SNVs in UTS2R can increase the sensitivity to remdesivir. Thus, this study provides mechanistic insights into remdesivir-mediated cardiac side effects and the therapeutic opportunity to prevent the aversive events in the future.

## MATERIALS AND METHODS

### Study design

The overall goal of this study was to explore the molecular mechanisms of the remdesivir-related cardiotoxicity in anti-COVID-19 therapy. We first performed the GPCR screening using major anti-COVID-19 drugs as ligands (n =3 per GPCR). We validated the results of GPCR screening by concentration-response analysis and determined the EC_50_ and E_max_ values of remdesivir against UTS2R (n =16). We also performed docking simulation using AlphaFold and determined the key amino acid residues that were essential for remdesivir-UTS2R binding as demonstrated by a mutagenesis study (n ≥3). We next examined the cardiotoxic potential of remdesivir. Using hiPS-derived cardiomyocytes, we found that the proarrhythmic risk of remdesivir is largely mediated by UTS2R (n=3). Moreover, using NRCMs, we showed that the decrease in contraction force by remdesivir is mediated by the UTS2R-Gα_i/o_ axis (n=3). Finally, the effects of SNVs in the *UTS2R* gene on remdesivir-mediated receptor activation was evaluated using GPCR assay (n =3 per SNV).

### Chemicals

Remdesivir (GS-5734), GS-441524, and sofosbuvir (GS-7977) were purchased from Selleck Chemicals. Favipiravir was purchased from Tokyo Chemical Industry. Fibronectin was purchased from Sigma-Aldrich. All other reagents were of analytical grade and obtained from commercial sources.

### SNV in the human *UTS2R* coding region

To search for SNVs in *UTS2R*, we used a public database of 14,000 healthy Japanese individuals, called 14KJPN, released by Tohoku University’s Tohoku Medical Megabank Organization (ToMMo, https://jmorp.megabank.tohoku.ac.jp/)(38), which covers variant frequency data.

### TGFα-shedding assay-based GPCR screening

To measure the activation of the GPCR, a transforming growth factor-α (TGFα) shedding assay was performed as described previously(*26*). Briefly, HEK293A cells were seeded in a 12-well culture plate. A total of 348 pCAG plasmids encoding an untagged GPCR including UTS2R construct were prepared (Table S1). The UTS2R mutants, including 110 missense mutants corresponding to the human SNVs in the *UTS2R* gene, were generated by introducing single-point mutations using a KOD-Plus-Mutagenesis kit. Cells were transfected using a polyethylenimine (PEI) reagent (2.5 µl of 1 mg/ml per well hereafter; Polysciences) with these UTS2R plasmids (100 ng per well hereafter), together with the plasmids encoding alkaline phosphatase (AP)-tagged TGFα (AP-TGFα; 250 ng). UTS2R activation assays were performed with endogenous G protein unless otherwise specified. Chimeric Gα subunit proteins (mixture of Gα_q/s_, Gα_q/i1_, Gα_q/i3_, Gα_q/o_, Gα_q/z_, Gα_q/12_, Gα_q/13_, and Gα_16_; each 10 ng) were used for initial screening of 348 GPCRs (Fig. 1a). After a 24-h culture, the transfected cells were harvested and collected by centrifugation. Cells were suspended in Hank’s Balanced Salt Solution (HBSS) containing 5 mM HEPES (pH 7.4) and seeded in a 96-well plate. After a 30-min incubation, the test compound was added to the cells.

After 1-h incubation, the conditioned media were transferred to an empty 96-well plate. The AP reaction solution (a mixture of 10 mM *p*-nitrophenylphosphate (*p*-NPP), 120 mM Tris-HCl (pH 9.5), 40 mM NaCl, and 10 mM MgCl_2_) was added to plates containing cells and conditioned media. The absorbance at a wavelength of 405 nm was measured using a microplate reader (Molecular Devices) before and after a 1-h incubation of the plates at room temperature. AP-TGFα release was calculated as described previously(*26*). The AP-TGFα release percentages were fitted to a four-parameter sigmoidal concentration-response curve, using Prism 9 software (GraphPad Prism), and the EC_50_ and E_max_ values were obtained.

### β-arrestin recruitment assay

NanoBiT enzyme complementation-based β-arrestin recruitment assay was performed as described previously (*54*). Briefly, HEK293A cells were seeded in a 6-well culture plate and transfected using a PEI reagent (5 µl of 1 mg/ml per well hereafter) with a mixture of plasmids consisting of 500 ng ssHA-FLAG-UTS2R-SmBiT construct (N-terminal hemagglutinin signal sequence followed by an FLAG epitope tag, plus C-terminal SmBiT with a 15-amino acid flexible linker), 100 ng LgBiT-β-arrestin construct (N-terminal LgBiT with a 15-amino acid flexible linker) and 400 ng empty plasmid. After a 24-hour culture, the transfected cells were harvested, suspended in 2 ml of assay buffer (Hanks’ Balanced Salt Solution containing 5 mM HEPES (pH 7.4) and 0.01% bovine serum albumin) and seeded in a 96-well plate at a volume of 80 µl per well. Coelenterazine (20 µl of 50 µM diluted in the assay buffer) was added to the cell plates followed by incubation at room temperature for 2 h in the dark. After measurement of baseline luminescence using a Spectra Max L Microplate Reader (Molecular Devices), remdesivir or UT2 (20 µl of 6X concentration) was added. Luminescence was measured every 20 s after compound addition. The average luminescent signal over 5–10 min was normalized to an initial value. The fold-change values were further normalized to that of the vehicle-treated condition and were fitted to a four-parameter sigmoidal concentration-response curve to obtain EC_50_ values using Prism 9 software.

### Docking simulation

A human UTS2R structure was prepared from the AlphaFold Protein Structure Database (*55, 56*) (AlphaFold Monomer v2.0) by trimming the lid-like N-terminus region (1–42) with a low pLDDT score. Hydrogen atoms were added to the trimmed receptor with the program Reduce (*57*). Remdesivir was docked in the orthosteric pocket of the receptor by AutoDockFR (*58*) with 50 genetic algorithm evolutions and a maximum of 2,500,000 evaluations. Docking poses were clustered at the 2 Å cutoff, and the top five clusters were inspected based on the criterion of hydrogen bonding between the receptor and remdesivir. Structures were visualized and analyzed with CueMol and PyMOL (Schrödinger, Inc.).

### Western blot analysis

For blots of pERK, ERK, pAKT, AKT, and β-actin, UTS2R-transfected and serum-starved HEK293A cells were stimulated with 0.1 or 1 μM remdesivir or 10 or 100 nM UT2. Before stimulation with these compounds, cells were incubated with urantide (Peptide Institute) for 30 min at 100 nM (for UTS2R inhibition), or with 150 ng/mL pertussis toxin (PTX; Wako) overnight (for Gα_i/o_ inhibition). For blots of mitochondrial proteins (Total OXPHOS, MT-COI, TFAM, and VDAC), 1 or 10 μM remdesivir or 1 or 10 μM GS-441524 was added to UTS2R-transfected HEK293A cells for 48 h. Whole-cell lysates were prepared in RIPA Lysis Buffer (Thermo Fischer Scientific) containing protease inhibitors (Thermo Fischer Scientific) and phosphatase inhibitors (Nacalai Tesque). Cell lysates were sonicated and clarified by centrifugation. Protein concentrations were measured using the BCA Protein Assay Kit (Pierce). Samples were separated using SDS polyacrylamide gels and transferred onto polyvinylidene difluoride (PVDF) membranes. The blots were blocked with 5% non-fat milk in TBS-T (Tris-buffered saline with 0.1% Tween-20) for 1 h, then probed using various primary antibodies. The primary antibodies and working dilutions used were as follows: pERK (AB_331646), 1:2000; ERK (AB_330744), 1:2000; pAKT (AB_329825), 1:1000; AKT (AB_329827), 1:1000; βactin (AB_10697039), 1:10000; total OXPHOS (AB_2756818), 1:10000; MTCO1 (AB_2084810), 1:2000; TFAM (AB_10841294), 1:1000; VDAC (AB_2272627), 1:1000. Target antigens were incubated with appropriate HRP-conjugated secondary antibodies (Cell Signaling) and were visualized by an ECL substrate (GE Healthcare).

### Quantification of UTS2R mRNA in tissues

To assess the human UTS2R (hUTS2R) mRNA expression in normal adult human tissues, commercially available cDNAs originating from various human tissues were purchased from TaKaRa (detailed sample information is given in Table S3) and 2 ng of cDNA were subjected to quantitative PCR (qPCR) analysis (Rotor-Gene Q, Qiagen), as recommended by the manufacturers. To assess mouse UTS2R (mUTS2R) mRNA expression in normal adult mouse tissues, C57BL/6J male mice were purchased from Japan Clea, and RNA was isolated from multiple tissues. RNA samples were reverse-transcribed into cDNA and were subjected to qPCR analysis. The primers used for the relative quantification are shown in Table S4. The expression levels of hUTS2R and mUTS2R were normalized to housekeeping genes of G3PDH and 18s rRNA, respectively.

### Cell culture of human induced pluripotent stem cell-derived cardiomyocytes

Human induced pluripotent stem cell-derived cardiomyocytes (hiPSC-CMs) were purchased from Fujifilm Cellular Dynamics, Inc. (CDI; iCell cardiomyocytes 2.0). To prepare the probe (MED- PG515A, Alpha MED Sciences), the recording areas were coated with fibronectin and dissolved in Dulbecco’s phosphate-buffered saline to create a 50 µg/mL solution. The recording area was covered with 1-2 µL fibronectin solution, and the probes were incubated at 37°C for at least 1 h. Cells were thawed and suspended in iCell Cardiomyocytes Plating Medium (CDI) and plated onto the probes at a density of 3.5×10^4^ cells in 2 µL plating medium. The cells were incubated at 37°C in 5% CO_2_ for 3–4 h before filling each probe with 1 mL iCell cardiomyocytes maintenance medium (CDI), which was used as the culture medium. The medium was changed every 2–3 days, with the cells cultured in the probes for 5–14 days to obtain a sheet of cardiomyocytes with spontaneous and synchronous electrical activity.

### Field potential recordings and data analysis

Field potential (FP) recording was carried out as previously described with slight modifications (*34, 59*). Briefly, before measuring FPs, iCell cardiomyocytes sheets were equilibrated for at least 4 h in a fresh culture medium using a 5% CO_2_ incubator at 37°C. After equilibration, the probes were transferred to the multi-electrode array (MEA) system (MED64, Alpha med Scientific) and incubated in a humidified 5% CO_2_ atmosphere. The stability and constancy of the waveforms were confirmed by monitoring the signals for at least 30 min. Once the FPs reached a stable state, they were recorded for 10 min at the baseline and then 24, 48, and 72 h after drug treatment. The stock solutions of drugs were prepared in dimethyl sulfoxide (DMSO) or distilled water at a 1000-fold target concentration. Stock solution was diluted in a culture medium to a final concentration of 0.1% DMSO.

FP data analysis was conducted, as described previously (*34*). Briefly, the FP duration (FPD) was defined as the duration from the 1^st^ to the 2^nd^ peak in FP. The FPD was corrected for the beat rate (inter-spike interval (ISI)) with Fridericia’s formula, which was the primary method of correction in this study (FPDcF = FPD/ (ISI/1000)^1/3^). The ISI and FPD value were averaged from the last 30 beats or 30 beats at time points exhibiting a stable ISI and FPD.

### Isolation of neonatal rat cardiomyocyte

Three-day-old Sprague–Dawley rat pups were purchased from Japan SLC Inc. for the isolation of neonatal rat cardiomyocytes (NRCMs), performed as previously described (*60*).

### Observation and analysis of contractility with pacing

NRCMs (4×10^5^ cells/well) were seeded on 3.5-cm glass-bottomed dishes in serum-free DMEM. Cells were treated with remdesivir (1 μM) for 48 hours at 37 °C and 5%CO_2_. Urantide (100 nM) was added at the same time. Pertussis toxin (PTX; 150 ng/mL) was treated 18 h before adding remdesivir, and YM-254890 (1 μM) was treated 30 minutes before remdesivir addition. Microscopic images of NRCMs were recorded for 13 seconds under pacing conditions. The pacing of NRCM was stimulated at 4 V every second, with a frequency of 10 msec, using an electrical stimulator (NIHON KOHDEN, SEN-3301). Imaging data were acquired at six frames per second using a BZ-X800 microscope (Keyence) and analyzed using Fiji software, as previously described (*61*).

### Statistical analysis

Statistical analyses were performed with Prism Software (GraphPad) and methods are described in the figure legends. Symbols are mean values and error bars denote standard error of the mean (SEM). Representative results from at least three independent experiments are shown for every figure, unless stated otherwise in the figure legends.

## Acknowledgments

We are grateful for suggestions and technical support from all members of the Department of Modomics Biology & Medicine, Tohoku University. We thank Y. Takahata, H. Miyamoto for technical assistance, and Natalie D. DeWitt for critical reading and language editing.

## Funding

JSPS KAKENHI Grant Number JP20K18371, JP22H04628 and JP22H02813 (AO)

JSPS KAKENHI Grant Number JP21H05269 (MN)

JSPS KAKENHI Grant Number JP21H04791 and JP21H051130 (AI)

JSPS KAKENHI Grant Number JP21H02659 and JP21H05265 (FYW)

A Grant-in-Aid for Scientific Research from the Ministry of Education, Culture, Sports, Science, and Technology, Japan 21H02634 (YK [Kanda])

The Research on Regulatory Harmonization and Evaluation of Pharmaceuticals, Medical Devices, Regenerative and Cellular Therapy Products, Gene Therapy Products, and Cosmetics from Japan Agency for Medical Research and Development (AMED)

JP21mk0101189 (YK [Kanda])

AMED BINDS JP22ama121031(MN)

AMED LEAP JP20gm0010004 (AI)

AMED BINDS JP20am0101095 (AI)

AMED 21he2302008j0002 (FYW)

FOREST JPMJFR215T from Japan Science and Technology Agency (JST) (AI)

JST Moonshot R&D JPMJMS2023 from JST (AI)

FOREST JPMJFR205Y from JST (FYW)

ERATO JPMJER2002 from JST (FYW)

Daiichi Sankyo Foundation of Life Science (AI)

Takeda Science Foundation (AI)

Uehara Memorial Foundation (AO)

Uehara Memorial Foundation (AI)

Inamori Foundation (AO)

Kowa Life Science Foundation (AO)

Mochida Memorial Foundation (AO)

Noguchi-Shitagau Research Grant (AO)

Japan Foundation for Applied Enzymology (AO)

The Mitsubishi Foundation (AO)

Terumo Life Science Foundation (AO)

Gout and uric acid foundation (AO)

The NOVARTIS Foundation (AO)

The Naito Foundation (AO)

Astellas Foundation for Research on Metabolic Disorders (AO)

## Author contributions

Conceptualization: AO, AI, FYW

Methodology: AO, TI, YK[Kato], SY, YK[Kanda], MN, AI

Investigation: AO, SO, TI, YK[Kato], SY, YI, AI

Visualization: AO, TI, YK[Kato], SY, AI, FYW

Funding acquisition: AO, YK[Kanda], MN, AI, FYW

Project administration: AI, FYW

Supervision: YK[Kanda], MN, AI, FYW

Writing – original draft: AO, TI, YK[Kato], SY, AI

Writing – review & editing: YK[Kanda], MN, AI, FYW

## Data and materials availability

All data are available in the main text or the supplementary materials.

---

**Fig. S1.**
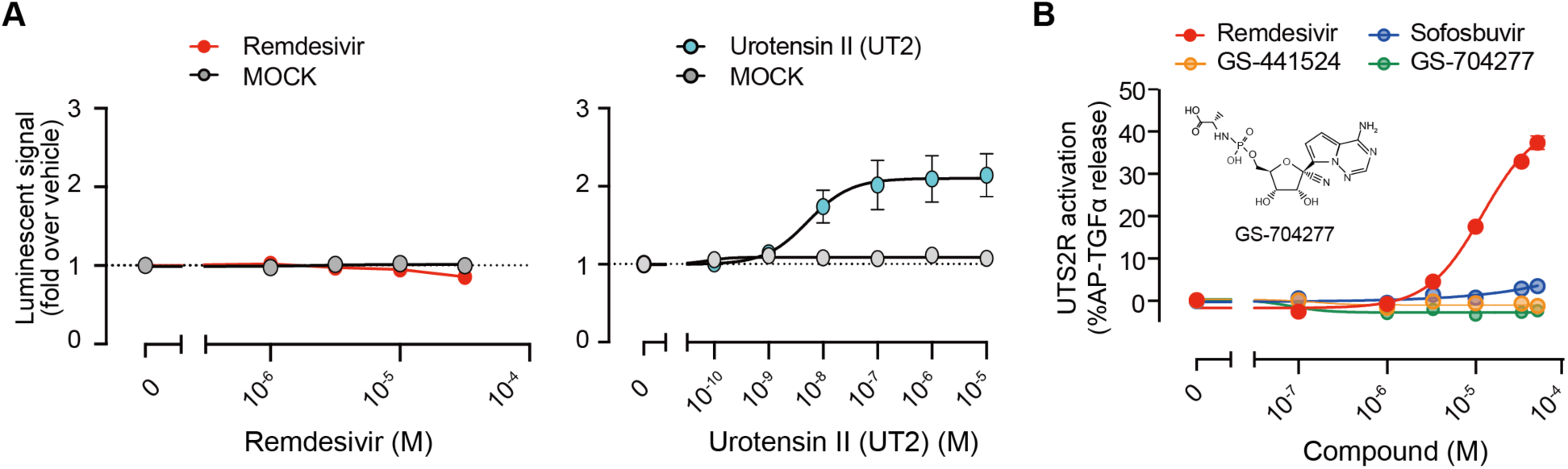
Activation of urotensin-II receptor (UTS2R) by remdesivir. **(A)** (left) Remdesivir- or (right) urotensin-II (UT2)-mediated β-arrestin 2 recruitment assay for UTS2R. **(B)** Remdesivir-related compounds-mediated TGFα-shedding response curves for UTS2R. Data are shown as means ± SEM (n ≥3).

**Fig. S2.**
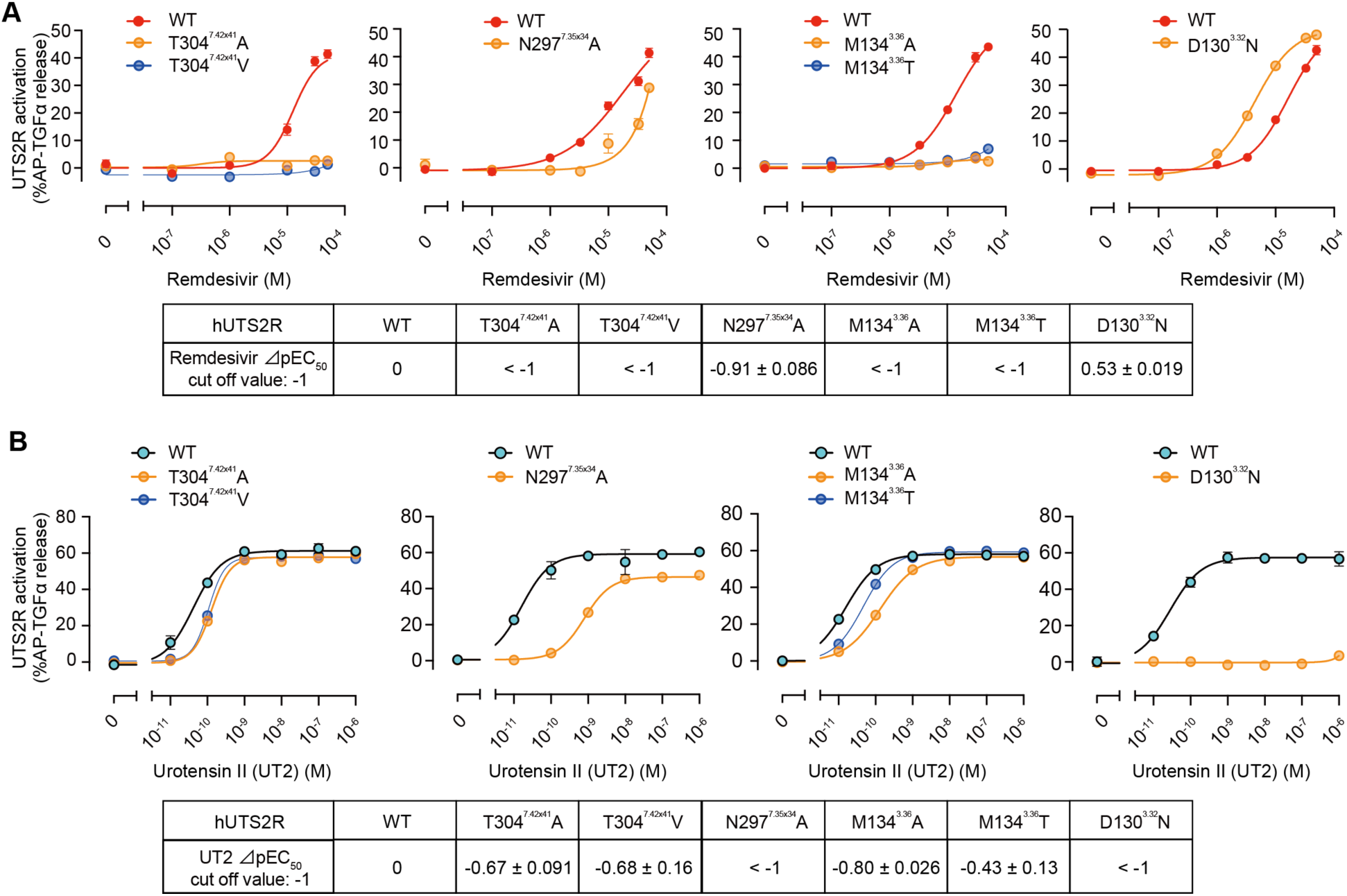
Effects of mutations on UTS2R activation. (A, B) Remdesivir (A)- and UT2 (B)- mediated TGFα-shedding response curves for UTS2R WT and indicated mutants.⊿pEC_50_ value = pEC_50_ mutant - pEC_50_ WT. The ⊿pEC_50_ cutoff value was set to -1. EC_50_ values were determined by the TGFα-shedding assay. Data are shown as means ± SEM (n ≥3).

**Fig. S3.**
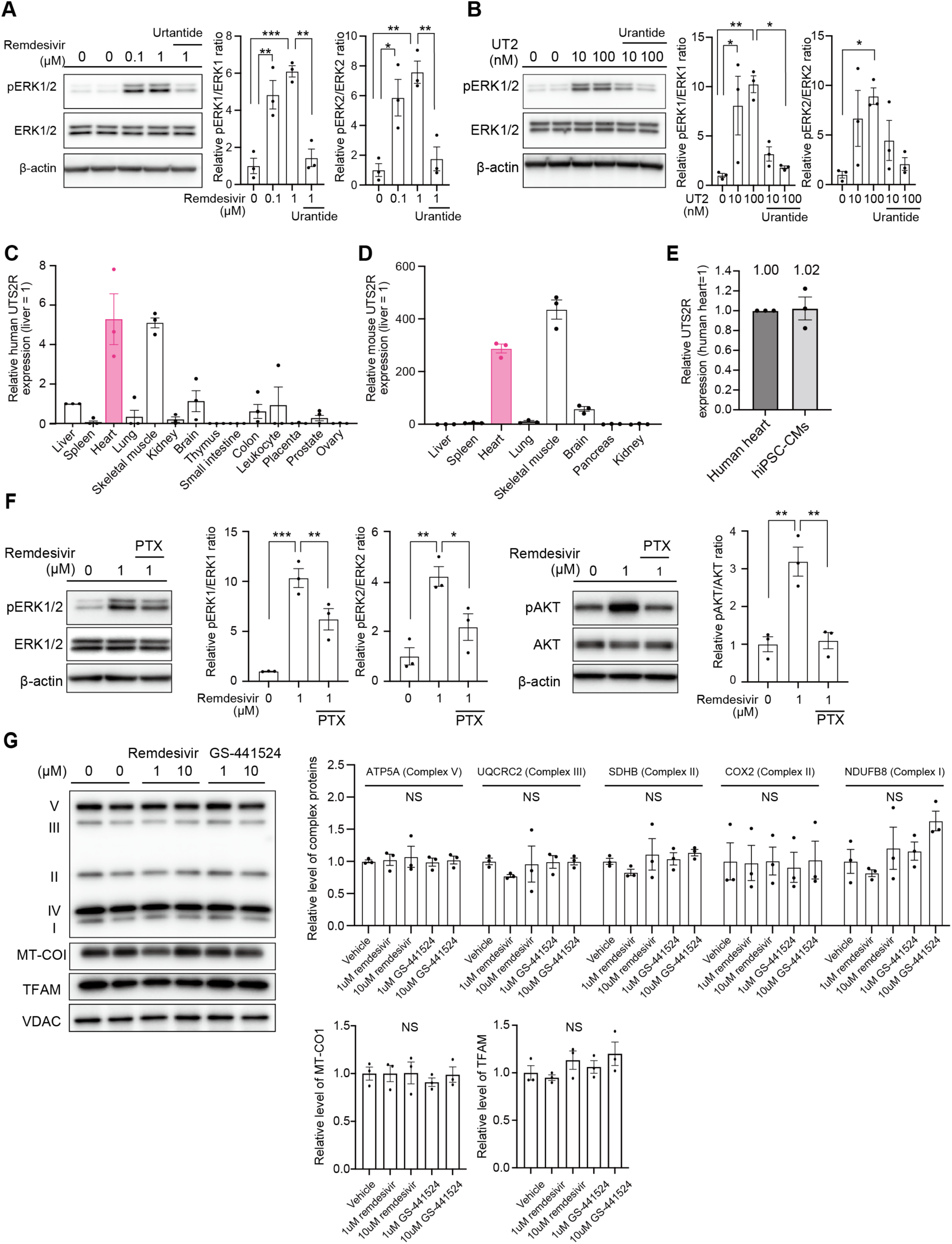
Intracellular signaling evoked by remdesivir-mediated UTS2R activation. **(A)** Serum-starved HEK293 cells overexpressing UTS2R were stimulated with the indicated concentrations of remdesivir for 72 h with or without urantide, a UTS2R antagonist, and the lysates subjected to western blotting analysis. ERK1 and ERK2 activation ratio (pERK1/ERK1 and pERK2/ERK2) were calculated with data normalized to the vehicle. * p <0.05, ** p <0.01, *** p <0.001 by Tukey’s multiple comparisons test. **(B)** Serum-starved HEK293 cells overexpressing UTS2R were stimulated with the indicated concentrations of UT2 for 5 min with or without urantide, a UTS2R antagonist, and the lysates subjected to western blotting analysis. ERK1 and ERK2 activation ratio were calculated with data normalized to the vehicle. * p <0.05, ** p <0.01 by Tukey’s multiple comparisons test. **(C)** Relative expression levels of UTS2R in normal human tissues were assessed by qPCR using the Human Multiple Tissue cDNA (MTC) panel. **(D)** Relative expression levels of Uts2r in adult mouse tissues as assessed by qPCR. **(E)** Comparison of relative gene expression levels of UTS2R between human heart (using MTC panel) and human-induced pluripotent stem cell-derived cardiomyocytes (hiPSC-CMs). **(F)** Representative western blot for phosphorylation of (left) ERK1/2 and (right) protein kinase B (AKT). Serum-starved HEK293 cells overexpressing UTS2R were stimulated with the indicated concentrations of remdesivir for 48 h. For G_i/o_ protein inhibition, cells were incubated with pertussis toxin (PTX) for at least 18 h at 150 ng/mL. ERK1, ERK2, and AKT activation ratio were calculated with data normalized to the vehicle. * p <0.05, ** p <0.01, *** p <0.001 by Tukey’s multiple comparisons test. **(G)** Expression levels of mitochondrial respiratory complex proteins in cells treated with remdesivir or GS-441524. HEK293 cells overexpressing UTS2R were stimulated with the indicated concentrations of remdesivir or GS-4415424 for 48 h. Mitochondrial respiratory complex proteins were normalized on VDAC and calculated with data normalized to the vehicle by Tukey’s multiple comparisons test

**Fig. S4.**
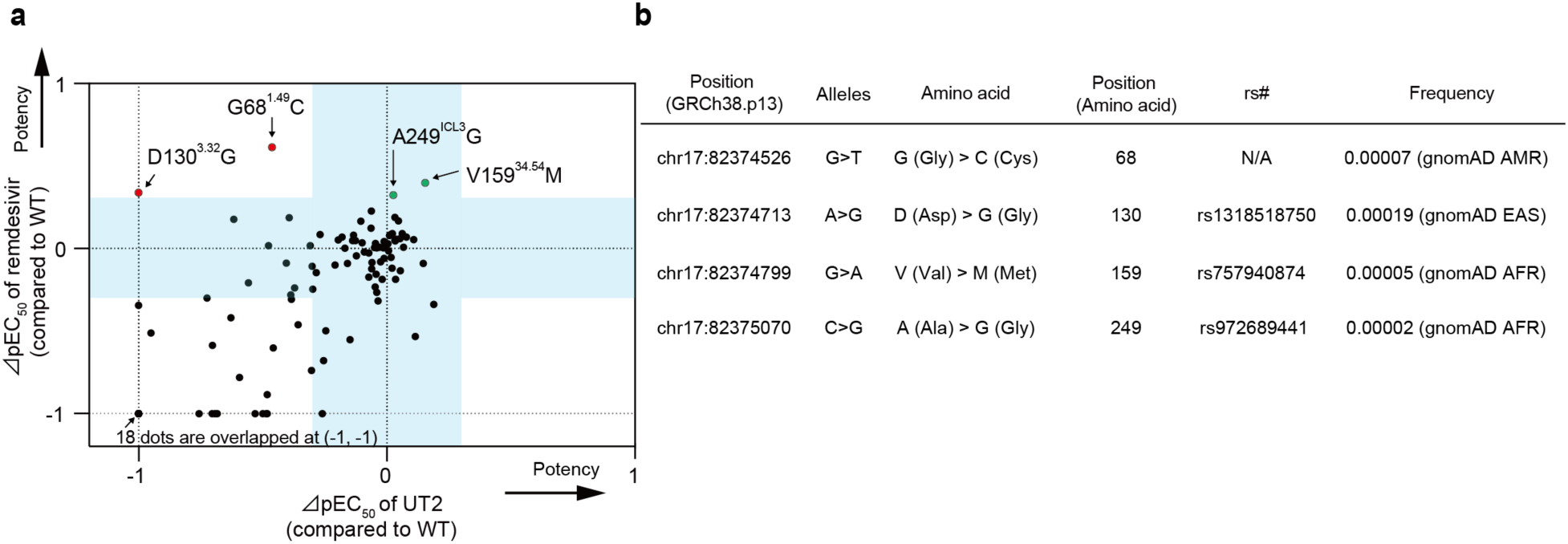
Effects of SNV on remdesivir-mediated UTS2R activation. **(A)** Scatter plot of ⊿pEC_50_ of remdesivir vs. ⊿pEC_50_ of UT2 for 110 missense human SNVs. Light blue bands represent the range of -0.3< ⊿pEC_50_ <0.3, which corresponds to the range of >0.5-fold and <2-fold change in EC_50_ compared to the WT receptor. Mutations that increase the receptor sensitivity toward remdesivir (⊿pEC_50_ >0.3) but not toward the UT2 (⊿pEC_50_ <0) are marked in red. Mutations that increase the receptor sensitivity toward both remdesivir and UT2 (⊿pEC_50_ >0.3 for remdesivir, 0< ⊿pEC_50_ <0.3 for UT2) are marked in green. **(B)** Allele frequencies of the four gain-of-function mutations (G68^1.49^C, D130^3.32^G, V159^34.54^M, and A249^ICL3^G). Data are modified from the 14KJPN Genome Reference Panel (https://jmorp.megabank.tohoku.ac.jp/202112/).

**Data file S1. TGF-α shedding assay screening data**

**Data file S2. Effects of missense SNVs in UTS2R gene on receptor activation Data file S3. Detailed sample information of pooled human cDNA**

**Data file S4. Sequences of primers used for quantitative PCR**

## References

1. K. L. Seley-Radtke, M. K. Yates, The evolution of nucleoside analogue antivirals: A review for chemists and non-chemists. Part 1: Early structural modifications to the nucleoside scaffold. Antiviral Res 154, 66–86 (2018).

2. M. Chien et al., Nucleotide Analogues as Inhibitors of SARS-CoV-2 Polymerase, a Key Drug Target for COVID-19. J Proteome Res 19, 4690–4697 (2020).

3. A. S. Alanazi, E. James, Y. Mehellou, The ProTide Prodrug Technology: Where Next? ACS Med Chem Lett 10, 2–5 (2019).

4. T. P. Sheahan, et al., Broad-spectrum antiviral GS-5734 inhibits both epidemic and zoonotic coronaviruses. Sci Transl Med 9, (2017).

5. D. Siegel et al., Discovery and Synthesis of a Phosphoramidate Prodrug of a Pyrrolo[2,1- f][triazin-4-amino] Adenine C-Nucleoside (GS-5734) for the Treatment of Ebola and Emerging Viruses. J Med Chem 60, 1648–1661 (2017).

6. T. K. Warren et al., Therapeutic efficacy of the small molecule GS-5734 against Ebola virus in rhesus monkeys. Nature 531, 381–385 (2016).

7. J. H. Beigel et al., Remdesivir for the Treatment of Covid-19 - Final Report. N Engl J Med 383, 1813–1826 (2020).

8. R. L. Gottlieb et al., Early Remdesivir to Prevent Progression to Severe Covid-19 in Outpatients. N Engl J Med 386, 305–315 (2022).

9. Coronavirus Disease 2019 (COVID-19) Treatment Guidelines. (2022).

10. S. Y. Jung et al., Cardiovascular events and safety outcomes associated with remdesivir using a World Health Organization international pharmacovigilance database. Clin Transl Sci 15, 501–513 (2022).

11. M. Haghjoo et al., Effect of COVID-19 medications on corrected QT interval and induction of torsade de pointes: Results of a multicenter national survey. Int J Clin Pract 75, e14182 (2021).

12. D. Liu et al., Adverse Cardiovascular Effects of Anti-COVID-19 Drugs. Front Pharmacol 12, 699949 (2021).

13. A. K. Gupta, B. M. Parker, V. Priyadarshi, J. Parker, Cardiac Adverse Events With Remdesivir in COVID-19 Infection. Cureus 12, e11132 (2020).

14. W. J. Hu et al., Pharmacokinetics and tissue distribution of remdesivir and its metabolites nucleotide monophosphate, nucleotide triphosphate, and nucleoside in mice. Acta Pharmacol Sin 42, 1195–1200 (2021).

15. C. J. Gordon, E. P. Tchesnokov, R. F. Schinazi, M. Götte, Molnupiravir promotes SARS-CoV-2 mutagenesis via the RNA template. J Biol Chem 297, 100770 (2021).

16. A. Jayk Bernal et al., Molnupiravir for Oral Treatment of Covid-19 in Nonhospitalized Patients. N Engl J Med 386, 509–520 (2022).

17. Y. Furuta et al., T-705 (favipiravir) and related compounds: Novel broad-spectrum inhibitors of RNA viral infections. Antiviral Res 82, 95–102 (2009).

18. A. K. Padhi, J. Dandapat, P. Saudagar, V. N. Uversky, T. Tripathi, Interface-based design of the favipiravir-binding site in SARS-CoV-2 RNA-dependent RNA polymerase reveals mutations conferring resistance to chain termination. FEBS Lett 595, 2366–2382 (2021).

19. W. Wen et al., Efficacy and safety of three new oral antiviral treatment (molnupiravir, fluvoxamine and Paxlovid) for COVID-19 : a meta-analysis. Ann Med 54, 516–523 (2022).

20. D. T. Hung et al., The efficacy and adverse effects of favipiravir on patients with COVID-19: A systematic review and meta-analysis of published clinical trials and observational studies. Int J Infect Dis 120, 217–227 (2022).

21. K. A. Jacobson, Z. G. Gao, Adenosine receptors as therapeutic targets. Nat Rev Drug Discov 5, 247–264 (2006).

22. A. Ogawa et al., N(6)-methyladenosine (m(6)A) is an endogenous A3 adenosine receptor ligand. Mol Cell 81, 659–674.e657 (2021).

23. G. Rengo, A. Lymperopoulos, W. J. Koch, Future g protein-coupled receptor targets for treatment of heart failure. Curr Treat Options Cardiovasc Med 11, 328–338 (2009).

24. M. Sato, Y. Ishikawa, Accessory proteins for heterotrimeric G-protein: Implication in the cardiovascular system. Pathophysiology 17, 89–99 (2010).

25. D. Wacker, R. C. Stevens, B. L. Roth, How Ligands Illuminate GPCR Molecular Pharmacology. Cell 170, 414–427 (2017).

26. A. Inoue et al., TGFα shedding assay: an accurate and versatile method for detecting GPCR activation. Nat Methods 9, 1021–1029 (2012).

27. Summary-compassionate-use-remdesivir-gilead_en.pdf. (2020).

28. H. Castel et al., The G Protein-Coupled Receptor UT of the Neuropeptide Urotensin II Displays Structural and Functional Chemokine Features. Front Endocrinol (Lausanne) 8, 76 (2017).

29. J. A. Ballesteros, H. Weinstein, Integrated methods for the construction of three-dimensional models and computational probing of structure-function relations in G protein-coupled receptors. Methods in Neurosciences 25, 366–428 (1995).

30. H. S. Biswal, E. Gloaguen, Y. Loquais, B. Tardivel, M. Mons, Strength of NH···S Hydrogen Bonds in Methionine Residues Revealed by Gas-Phase IR/UV Spectroscopy. J Phys Chem Lett 3, 755–759 (2012).

31. P. R. Pokkuluri et al., Factors contributing to decreased protein stability when aspartic acid residues are in beta-sheet regions. Protein Sci 11, 1687–1694 (2002).

32. J. Pereira-Castro, C. Brás-Silva, A. P. Fontes-Sousa, Novel insights into the role of urotensin II in cardiovascular disease. Drug Discov Today 24, 2170–2180 (2019).

33. C. L. Mummery et al., Differentiation of human embryonic stem cells and induced pluripotent stem cells to cardiomyocytes: a methods overview. Circ Res 111, 344–358 (2012).

34. S. Yanagida, A. Satsuka, S. Hayashi, A. Ono, Y. Kanda, Comprehensive Cardiotoxicity Assessment of COVID-19 Treatments Using Human-Induced Pluripotent Stem Cell-Derived Cardiomyocytes. Toxicol Sci 183, 227–239 (2021).

35. L. X. Cubeddu, QT prolongation and fatal arrhythmias: a review of clinical implications and effects of drugs. Am J Ther 10, 452–457 (2003).

36. N. Wettschureck, S. Offermanns, Mammalian G proteins and their cell type specific functions. Physiol Rev 85, 1159–1204 (2005).

37. M. Nishida et al., G alpha(i) and G alpha(o) are target proteins of reactive oxygen species. Nature 408, 492–495 (2000).

38. S. Tadaka, et al., jMorp updates in 2020: large enhancement of multi-omics data resources on the general Japanese population. Nucleic Acids Res 49, D536–d544 (2021).

39. K. E. Lasser et al., Timing of new black box warnings and withdrawals for prescription medications. Jama 287, 2215–2220 (2002).

40. V. Michaud et al., Risk Assessment of Drug-Induced Long QT Syndrome for Some COVID-19 Repurposed Drugs. Clin Transl Sci 14, 20–28 (2021).

41. M. Nabati, H. Parsaee, Potential Cardiotoxic Effects of Remdesivir on Cardiovascular System: A Literature Review. Cardiovasc Toxicol 22, 268–272 (2022).

42. L. B. Day, H. Abdel-Qadir, M. Fralick, Bradycardia associated with remdesivir therapy for COVID-19 in a 59-year-old man. Cmaj 193, E612–e615 (2021).

43. M. I. Sanchez-Codez, M. Rodriguez-Gonzalez, I. Gutierrez-Rosa, Severe sinus bradycardia associated with Remdesivir in a child with severe SARS-CoV-2 infection. Eur J Pediatr 180, 1627 (2021).

44. S. A. Douglas, L. Tayara, E. H. Ohlstein, N. Halawa, A. Giaid, Congestive heart failure and expression of myocardial urotensin II. Lancet 359, 1990–1997 (2002).

45. M. L. Adamsick et al., Remdesivir in Patients with Acute or Chronic Kidney Disease and COVID-19. J Am Soc Nephrol 31, 1384–1386 (2020).

46. P. A. Insel, C. M. Tang, I. Hahntow, M. C. Michel, Impact of GPCRs in clinical medicine: monogenic diseases, genetic variants and drug targets. Biochim Biophys Acta 1768, 994–1005 (2007).

47. R. J. Lefkowitz, S. K. Shenoy, Transduction of receptor signals by beta-arrestins. Science 308, 512–517 (2005).

48. G. Esposito et al., EGFR trans-activation by urotensin II receptor is mediated by β-arrestin recruitment and confers cardioprotection in pressure overload-induced cardiac hypertrophy. Basic Res Cardiol 106, 577–589 (2011).

49. T. Korhonen, S. L. Hänninen, P. Tavi, Model of excitation-contraction coupling of rat neonatal ventricular myocytes. Biophys J 96, 1189–1209 (2009).

50. L. Sala et al., MUSCLEMOTION: A Versatile Open Software Tool to Quantify Cardiomyocyte and Cardiac Muscle Contraction In Vitro and In Vivo. Circ Res 122, e5–e16 (2018).

51. M. C. Sanguinetti, M. Tristani-Firouzi, hERG potassium channels and cardiac arrhythmia. Nature 440, 463–469 (2006).

52. M. Kwok et al., Remdesivir induces persistent mitochondrial and structural damage in human induced pluripotent stem cell derived cardiomyocytes. Cardiovasc Res, (2021).

53. E. P. Tchesnokov, J. Y. Feng, D. P. Porter, M. Götte, Mechanism of Inhibition of Ebola Virus RNA-Dependent RNA Polymerase by Remdesivir. Viruses 11, (2019).

54. Y. Hisano et al., Lysolipid receptor cross-talk regulates lymphatic endothelial junctions in lymph nodes. J Exp Med 216, 1582–1598 (2019).

55. J. Jumper et al., Highly accurate protein structure prediction with AlphaFold. Nature 596, 583–589 (2021).

56. M. Varadi et al., AlphaFold Protein Structure Database: massively expanding the structural coverage of protein-sequence space with high-accuracy models. Nucleic Acids Res 50, D439–d444 (2022).

57. J. M. Word, S. C. Lovell, J. S. Richardson, D. C. Richardson, Asparagine and glutamine: using hydrogen atom contacts in the choice of side-chain amide orientation. J Mol Biol 285, 1735–1747 (1999).

58. P. A. Ravindranath, S. Forli, D. S. Goodsell, A. J. Olson, M. F. Sanner, AutoDockFR: Advances in Protein-Ligand Docking with Explicitly Specified Binding Site Flexibility. PLoS Comput Biol 11, e1004586 (2015).

59. H. Ando et al., A new paradigm for drug-induced torsadogenic risk assessment using human iPS cell-derived cardiomyocytes. J Pharmacol Toxicol Methods 84, 111–127 (2017).

60. N. Kitajima et al., TRPC3 positively regulates reactive oxygen species driving maladaptive cardiac remodeling. Sci Rep 6, 37001 (2016).

61. Y. Matsuda, K. Takahashi, H. Kamioka, K. Naruse, Human gingival fibroblast feeder cells promote maturation of induced pluripotent stem cells into cardiomyocytes. Biochem Biophys Res Commun 503, 1798–1804 (2018).

